# Exploring the onset of B_12_-based mutualisms using a recently evolved Chlamydomonas auxotroph and B_12_-producing bacteria

**DOI:** 10.1101/2022.01.04.474942

**Authors:** Freddy Bunbury, Evelyne Deery, Andrew Sayer, Vaibhav Bhardwaj, Ellen Harrison, Martin J. Warren, Alison G. Smith

**Author notes:** Corresponding author: Professor Alison Smith, Department of Plant Sciences, University of Cambridge, Downing Street, Cambridge, CB2 3EA, United Kingdom. Carnegie Institution for Science, Department of Plant Biology, Stanford, California 94305, USA.

## Abstract

Cobalamin (vitamin B_12_), is a cofactor for crucial metabolic reactions in multiple eukaryotic taxa, including major primary producers such as algae, and yet only prokaryotes can produce it. Many bacteria can colonise the algal phycosphere, forming stable communities that gain preferential access to exudates and in return provide compounds, such as B_12_. Extended coexistence can then drive gene loss, leading to greater algal-bacterial interdependence. In this study, we investigate how a recently evolved B_12_-dependent strain of *Chlamydomonas reinhardtii*, metE7, forms a mutualism with certain bacteria, including the rhizobium *Mesorhizobium loti* and even a strain of the gut bacterium *E. coli* engineered to produce cobalamin. Although metE7 was supported by B_12_ producers, its growth in co-culture was slower than the B_12_-independent wild-type, suggesting that high bacterial B_12_ provision may be necessary to favour B_12_ auxotrophs and their evolution. Moreover, we found that an *E. coli* strain that releases more B_12_ makes a better mutualistic partner, and although this trait may be more costly in isolation, greater B_12_ release provided an advantage in co-cultures. We hypothesise that, given the right conditions, bacteria that release more B_12_ may be selected for, particularly if they form close interactions with B_12_-dependent algae.

**Originality-Significance statement:** Microalgae are fundamental to the global carbon cycle, and yet despite being photosynthetic they often rely on other organisms for micronutrients. One of these micronutrients is vitamin B_12_ (cobalamin), which they receive from bacteria. Many environmental studies support the widespread role of B_12_ in algal-bacterial mutualisms, so here we wished to investigate how these mutualisms may arise evolutionarily by using an experimentally evolved B_12_-dependent alga and various B_12_-producers. A B_12_-producing rhizobium, *Mesorhizobium loti*, could stably support the B_12_-dependent *Chlamydomonas reinhardtii* metE7 strain, and vice versa, but nutrient supplementation increased growth of both species further. metE7 could also be supported by *E. coli* strains engineered to produce B_12_, and engineering a strain to release a higher proportion of B_12_ led to better algal growth, which increased bacterial growth in turn. We suggest that as B_12_-based mutualisms develop, increased B_12_ release may be selected for and therefore lead to more productive symbioses.

## Introduction

The study of interactions within microbial communities is garnering increased attention as researchers continue to uncover the extent of microbial interdependence (Gude and Taga, 2019; Gralka *et al*., 2020), which is frequently driven by nutrient exchange. These connections extend to microbes across different domains of life, such as between humans and gut bacteria (Round and Mazmanian, 2009), and plants and their companions in the rhizosphere, including rhizobial bacteria (Udvardi and Poole, 2013) and arbuscular mycorrhizal fungi (Chen *et al*., 2018). Similarly, photosynthetic algae often support a range of heterotrophic bacteria in their phycosphere, a region near the algal cell surface analogous to the rhizosphere (Krohn-Molt *et al*., 2017; Seymour *et al*., 2017; Kimbrel *et al*., 2019), in return receiving specific compounds such as growth factors (Seyedsayamdost *et al*., 2011) or vitamins (Croft *et al*., 2005). For example, the marine alga *Ostreococcus tauri* can support the bacterium *Dinoroseobacter shibae* in co-culture by providing photosynthate, niacin, biotin and p-aminobenzoic acid, and obtain cobalamin and thiamine in return (Cooper *et al*., 2019). Cobalamin (vitamin B_12_) is a structurally complex cobalt-containing corrinoid molecule that is of particular interest in algal-bacterial interactions because it is only made by prokaryotes and yet more than half of microalgae require it for growth (Croft *et al*., 2005; Tang *et al*., 2010). B_12_ transfer among species, therefore, has an important role in maintaining community structure and function (Heal *et al*., 2017; Gómez-Consarnau *et al*., 2018; Sharma *et al*., 2019).

Nutrient amendment experiments of aquatic ecosystems have revealed that B_12_ or B_12_-producers frequently limit phytoplankton growth (Bertrand *et al*., 2007; Koch *et al*., 2011; Paerl *et al*., 2015; Joglar *et al*., 2020; Barber-Lluch *et al*., 2021), and laboratory experiments have investigated the effect of B_12_ supply on phytoplankton physiology (Bertrand *et al*., 2012; Heal *et al*., 2019; Koch and Trimborn, 2019; Nef *et al*., 2019). Studies have revealed the effects of B_12_ on the methionine cycle, C1 metabolism, and cell growth and division in *Euglena gracilis (Shehata and Kempner, 1978; Carell and Seeger, 1980), Tisochrysis lutea (Nef et al., 2019), Thalassiosira pseudonana (Heal et al., 2019)*, and *Chlamydomonas reinhardtii (Bunbury et al., 2020)*. In these species, B_12_ acts as a cofactor for the enzyme methionine synthase (METH), although some, like *C. reinhardtii*, also encode a B_12_-independent isoform (METE). It is the presence or absence of this isoform that is the best determinant of algal B_12_ dependence (Helliwell, 2017), and while B_12_ dependence is widespread, its complex phylogenetic distribution points to multiple instances of METE gene loss (Helliwell *et al*., 2011; Ellis *et al*., 2017).

Although B_12_ release may simply coincide with bacterial cell death (Haines and Guillard, 1974; Droop, 2007; Grant *et al*., 2014), several studies have shown that stable mutualisms can form between B_12_-producing bacteria and B_12_-requiring algae (Kazamia *et al*., 2012; Cruz-López and Maske, 2016; Peaudecerf *et al*., 2018), and there is some evidence that B_12_ supply may be regulated (Grant et al. 2014). In this case, the exchange of B_12_ between algae and bacteria necessitates appropriate export and import systems for this compound. Whether, and if so how, B_12_ export is regulated by producers on the other hand, is not known (Gude and Taga, 2019). The BtuBFCD complex in Gram-negative bacteria or BtuFCD complex in Gram-positive bacteria are the best characterised prokaryotic systems for B_12_ uptake (Rodionov *et al*., 2003; Degnan, Barry, *et al*., 2014), and although there is speculation that many vitamin transporters may be bidirectional, this has not been confirmed for cobalamin (Romine *et al*., 2017). Much less is known in algae about the molecular machinery, although a protein involved in algal B_12_ uptake was identified in the diatoms *Thalassiosira pseudonana* and *Phaeodactylum tricornutum (Bertrand et al., 2012)*, and subsequently in *C. reinhardtii* (Sayer et al. submitted), and B_12_ uptake proteins are now predicted by homology to exist in multiple algal species.

Whatever the mechanism of B_12_ release, its presence in the environment by ‘providers’ that generate this nutrient pool paves the way for the evolution of ‘providers’ into ‘cheaters’, that is, organisms that benefit from but do not contribute to the nutrient pool (Lenski *et al*., 2012). The high frequency with which only partial B_12_ biosynthesis pathways are found in bacterial genomes is testament to the evolutionary benefit of dispensing with this metabolically expensive process (Shelton *et al*., 2019). The Black Queen Hypothesis (BQH) seeks to explain this process of gene loss that results in reductive evolution, and how the equilibrium in the ratio of providers to cheaters is dependent upon the cost of production and leakiness of the public good (Morris, 2015). Leakiness is generally considered to be detrimental but unavoidable. However, in the context of a mutualistic cross-feeding relationship in an environment with a high spatial structure, increased leakiness may prove advantageous (Stump *et al*., 2018). A hypothesis can therefore be developed whereby it is only when B_12_-producing bacteria are closely associated with B_12_-requirers such as photosynthetic algae that provide something in return, that B_12_ release is evolutionarily favoured.

The unicellular chlorophyte, *C. reinhardtii*, is widely used as a model organism for researching photosynthesis and abiotic stress responses, (Sasso *et al*., 2018; Salomé and Merchant, 2019), but it has also been used to study microbial symbiosis (Hom and Murray, 2014; Calatrava *et al*., 2018; Durán *et al*., 2021). Here we use *C. reinhardtii* and B_12_-producing bacteria to investigate the inherent capacity for a simple mutualism without previous coevolution. We then investigate how the symbiotic dynamics differ between bacteria and the *C. reinhardtii* wild-type B_12_-independent strain versus its B_12_ dependent mutant as an analogy for the commensal-mutualism switch that would occur for algae that evolve B_12_ dependence. Perturbing the mutualistic co-cultures by nutrient addition reveals how each species responds to the presence of other members of a natural community and to what extent the interaction is regulated. Finally, using bacteria that release different amounts of B_12_, we investigate how increased B_12_ release, a disadvantage in axenic cultures, proves advantageous in the presence of B_12_-dependent algae.

## Results

### B_12_-dependent strain of *C. reinhardtii* takes up B_12_ produced by heterotrophic bacterium

*Chlamydomonas reinhardtii* is a model alga in part because it grows well on media containing only inorganic nutrients. However, Helliwell et al. (2015) were able to generate a vitamin B_12_-dependent mutant (hereafter metE7) by experimental evolution in the presence of high concentrations of B_12_. We wanted to test whether a B_12_-producing bacterium could support the growth of metE7 and whether this interaction would develop into anything more complex than passive nutrient cross-feeding. To identify a suitable B_12_-producer we first chose four soil bacteria (*Mesorhizobium loti* MAFF303099, *Sinorhizobium meliloti* RM1021, *Rhizobium leguminosarum* bv. viciae 3841 and *Pseudomonas putida*), which we hypothesised might co-occur in the environment with *C. reinhardtii* and could grow under similar conditions. We cultured the strains in minimal medium (TP) with glycerol in a 12 hr light-12 hr dark regime. After 6 days of culture, the amount of B_12_ in the media and cell fractions was determined (Figure S1). In the three rhizobia, the level of B_12_ was roughly equal in both fractions, indicating that a significant proportion of synthesised B_12_ is lost to the surroundings. In *P. putida*, however, which unlike the rhizobial strains encodes the outer membrane B_12_ transporter *btuB*, a substantially smaller portion of detectable B_12_ was in the media. Considering that the proportion of B_12_ in the medium was highest for *M. loti*, as well the fact that stable co-cultures form between *M. loti* and the *C. reinhardtii* relative, *Lobomonas rostrata* (Kazamia et al., 2012), we chose *M. loti* for further axenic and co-culture experiments.

Before establishing co-cultures, we quantified the effect of vitamin B_12_ on the growth of the *C. reinhardtii* mutant compared to both the ancestral strain from which it was derived and a revertant strain that was no longer dependent on exogenous cobalamin (see Experimental;(Helliwell *et al*., 2015)). The three strains were cultured for seven days with several B_12_ concentrations. As previously demonstrated (Bunbury *et al*., 2020), metE7 maximal cell density showed a clear dose-response to vitamin B_12_, while the growth of the ancestral and revertant strains (both containing a functional METE gene) was not increased by B_12_ (Figure 1A, Figure S2). Similarly, the effect of B_12_ supplementation on *M. loti* was quantified for both the wild-type strain and a B_12_ synthesis mutant (bluB^-^;(Shimoda *et al*., 2008)), which was confirmed to be unable to synthesise B_12_ (Figure S3B). Neither strain was affected by B_12_ supplementation.

**Figure 1.**
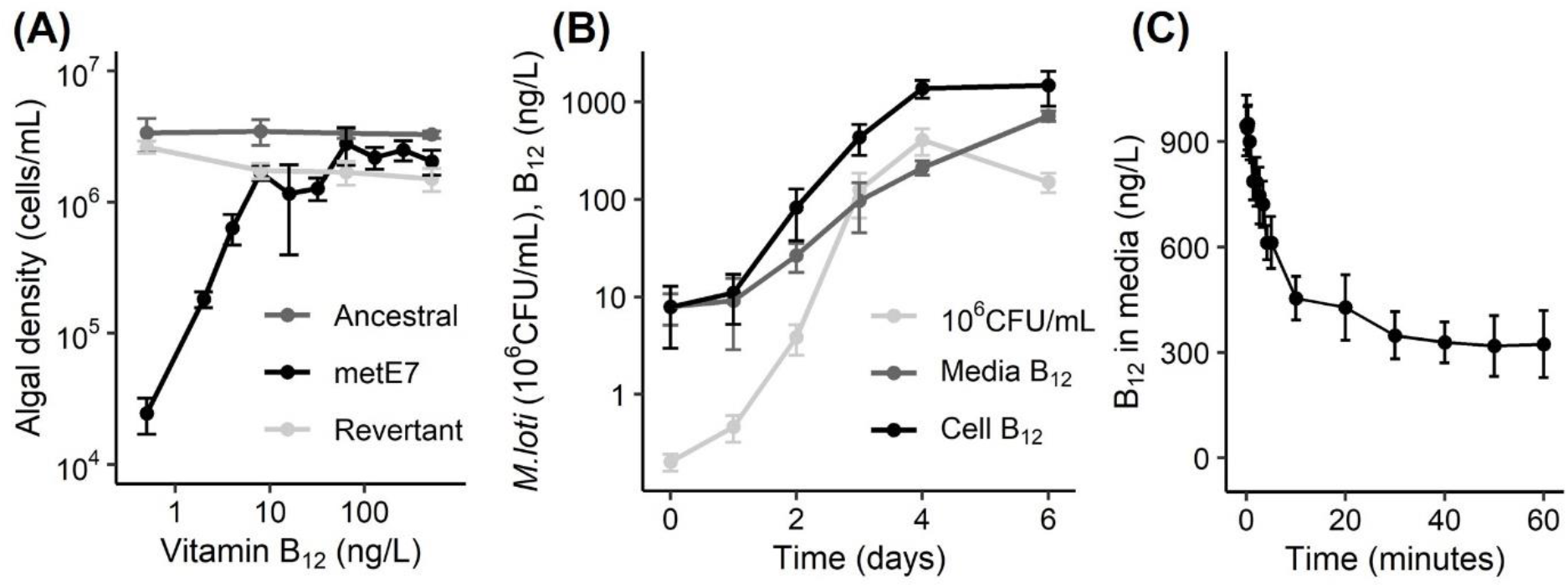
B_12_ dependent strain of *C. reinhardtii* takes up B_12_ produced by the heterotrophic bacterium *M. loti*. **(A)** The evolved metE7 mutant of *C. reinhardtii*, together with its ancestral line and a revertant (Helliwell et al. 2015), were cultured with a range of B_12_ concentrations in TP medium at 25 °C with constant illumination at 100 μE·m^-2^·s^-1^ for seven days, at which point the cell density in the cultures were measured. **(B)** *M. loti* was cultured in TP media with 0.1% glycerol at 25 °C with constant illumination at 100 μE·m^-2^·s^-1^ and measurements of cell density and B_12_ concentration were made over six days. **(C)** An *M. loti* culture which reached stationary phase was filtered through a 0.4 μm filter and metE7 cells starved of B_12_ were added to the filtrate at an OD730 nm of 0.1. This culture was then further filtered at multiple time intervals over the course of one hour to remove metE7 cells and the B_12_ concentration measured in this filtrate. Colours of points and lines are indicated in legends within the graphs, error bars represent standard deviations, n >= 3.

To study the growth and B_12_ production dynamics of *M. loti*, cultures were monitored over a 6-day period using the same conditions as for *C. reinhardtii*, but with 0.1% (v/v) added glycerol and no added B_12_. Figure 1B illustrates the rapid growth of *M. loti* by more than 1000-fold within four days followed by a small decline by day six. B_12_ concentration in the bacterial cell fraction increased but to a lesser extent than cell density, suggesting the amount of B_12_ per bacterium decreased, and the B_12_ in the media increased more slowly, but more consistently. To test whether this B_12_ could be used by metE7, we first completely removed all *M. loti* cells by filtering the *M. loti* culture through a 0.4 μm filter and adding the filtrate to a new syringe body. Axenic metE7 in late log-phase that had been precultured phototrophically with 200 ng·L^-1^ of B_12_ (an amount that is sufficient to avoid B_12_ deprivation while also preventing the accumulation of substantial B_12_ stores) was added to the *M. loti* filtrate at a final OD730 of 0.1. 1 ml filtered aliquots were then sampled at predetermined intervals over the next hour from this mixture of *M. loti* filtrate + metE7. Figure 1C shows that the concentration of B_12_ measured in the filtrate (that is B_12_ not taken up by metE7) declined substantially within 20 minutes from almost 1000 ng·L^-1^ to 400 ng·L^-1^ with little decline thereafter. This indicates that *M. loti* can produce and release significant quantities of B_12_ and this can be taken up rapidly by metE7. By contrast, we found *M. loti* is incapable of taking up exogenous B_12_ (Figure S3), and this may explain why B_12_ progressively accumulates over time in the media fraction of *M. loti* cultures.

### metE7 and *M. loti* support each other in mutualistic co-culture

After confirming metE7 could take up B_12_ produced by *M. loti*, we wanted to see how well *M. loti* would support metE7 in a co-culture with no B_12_ or exogenous organic carbon, so that *M. loti* would depend on the photosynthate provided by metE7. For comparison with this mutualistic interaction, we also set up two commensal co-cultures containing the ancestral or revertant strains (both B_12_-independent) with *M. loti*. Figure 2A shows that the ancestral and revertant strains were able to grow more quickly and to a higher density than metE7 from day 2 onwards, suggesting that low B_12_ levels were limiting metE7 growth. Despite the lower growth of metE7, *M. loti* density was similar in the commensal and mutualistic co-cultures (Figure 2B), but B_12_ concentrations were significantly lower in the mutualistic co-cultures up until the final day, when they equalised (Figure 2C, S4).

**Figure 2.**
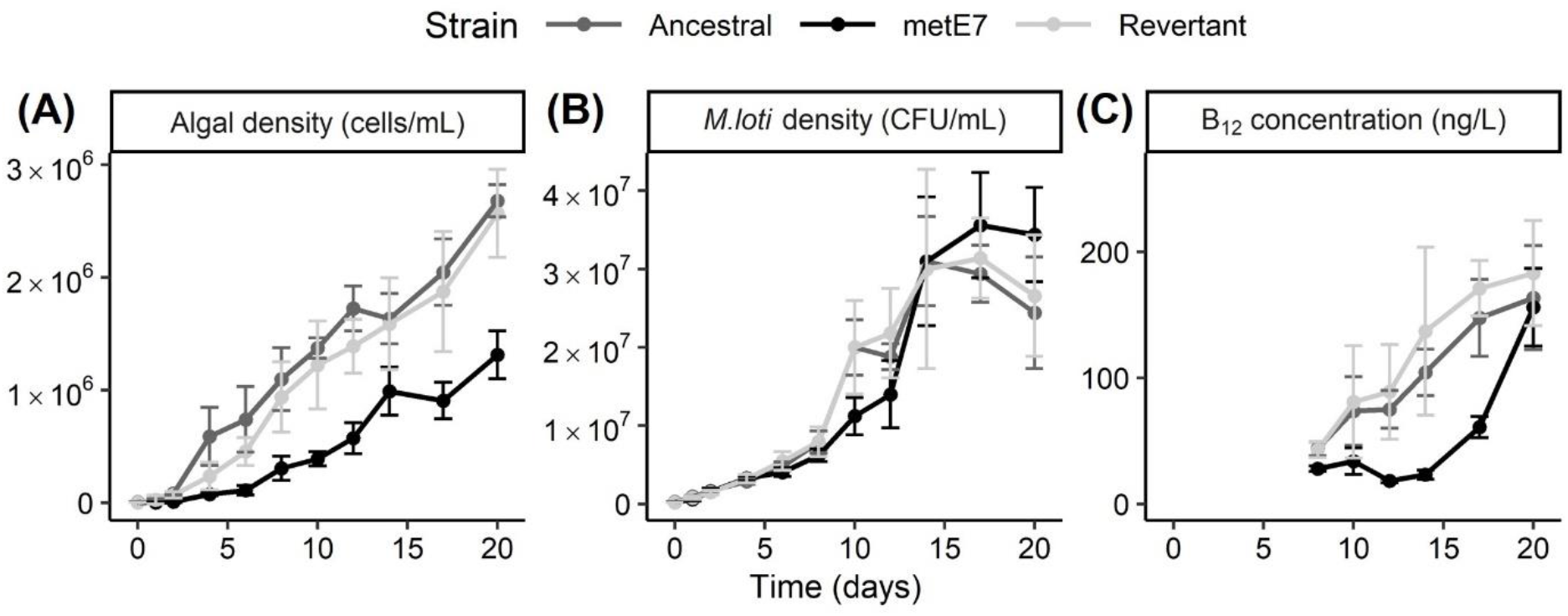
Comparison of commensal and mutualistic co-cultures between various strains of *C. reinhardtii* and *M. loti. M. loti* was co-cultured with the revertant or ancestral lines or metE7 in TP medium at 25 °C and with illumination at 100 μE·m^-2^·s^-1^ over a 16:8 hour light dark cycle for 20 days with periodic measurements of algal and bacterial density as well as B_12_ concentration **(A)** Measurement of algal density by particle counter reveals that ancestral and revertant line density increases at a faster rate than metE7 density in co-culture. **(B)** Counting colonies after plating serial dilutions of the cultures on TY media reveals that *M. loti* density is not significantly higher in co-culture with the ancestral or revertant lines than with metE7 on almost every day. **(C)** Total B_12_ concentration measured by *S. typhimurium* bioassay on aliquots of the co-cultures (media and cell fraction) reveals that B_12_ is generally higher in co-culture with the ancestral or revertant line. Grey = Ancestral line, black = metE7, error bars = standard deviation, n=5.

### metE7 and *M. loti* only partially support each other’s nutrient requirements and exhibit minimal partner regulation

Co-cultures provide some insight into how B_12_ producers and consumers naturally grow and survive in the environment, but ecosystems are clearly more complex due to the presence of multiple species and the fluctuations in physical conditions and nutrient availability. We chose to manipulate nutrient influx in the mutualistic co-cultures of metE7 and *M. loti* by testing the effect of added glycerol or B_12_. The co-cultures were initially grown phototrophically as before, with no B_12_ or glycerol, then split into three treatments: no nutrient addition, 200 ng·L^-1^ B_12_, or 0.02% (v/v) glycerol. The cultures were maintained semi-continuously for eight days by removing 10% of the culture for daily sampling and replacing it with the same volume of the respective media for each treatment.

As shown in Figure 3A, the metE7 density increased within one day of adding B_12_, but also increased in the cultures that were amended with glycerol, albeit with a delay of 1-2 days. metE7 densities in the glycerol and B_12_-supplemented cultures appeared to reach a new, roughly stable equilibrium level approximately 10 times higher than the control co-cultures. Glycerol addition increased *M. loti* levels by 10-fold relative to both the control and B_12_-supplemented cultures (Figure 3B). At no point did the *M. loti* density in the B_12_-supplemented culture significantly differ from the control suggesting that the increase in metE7 density did not translate into an increase in organic carbon available for *M. loti* growth. Total B_12_ increased significantly within one day of supplementation with either B_12_ itself or glycerol (Figure 3C), but only glycerol supplementation led to a substantial accumulation of B_12_ in the media (Figure S5A). Therefore, glycerol addition indirectly increased metE7 density, whereas B_12_-supplementation, which fuelled the growth of metE7, did not result in increased *M. loti* growth.

**Figure 3.**
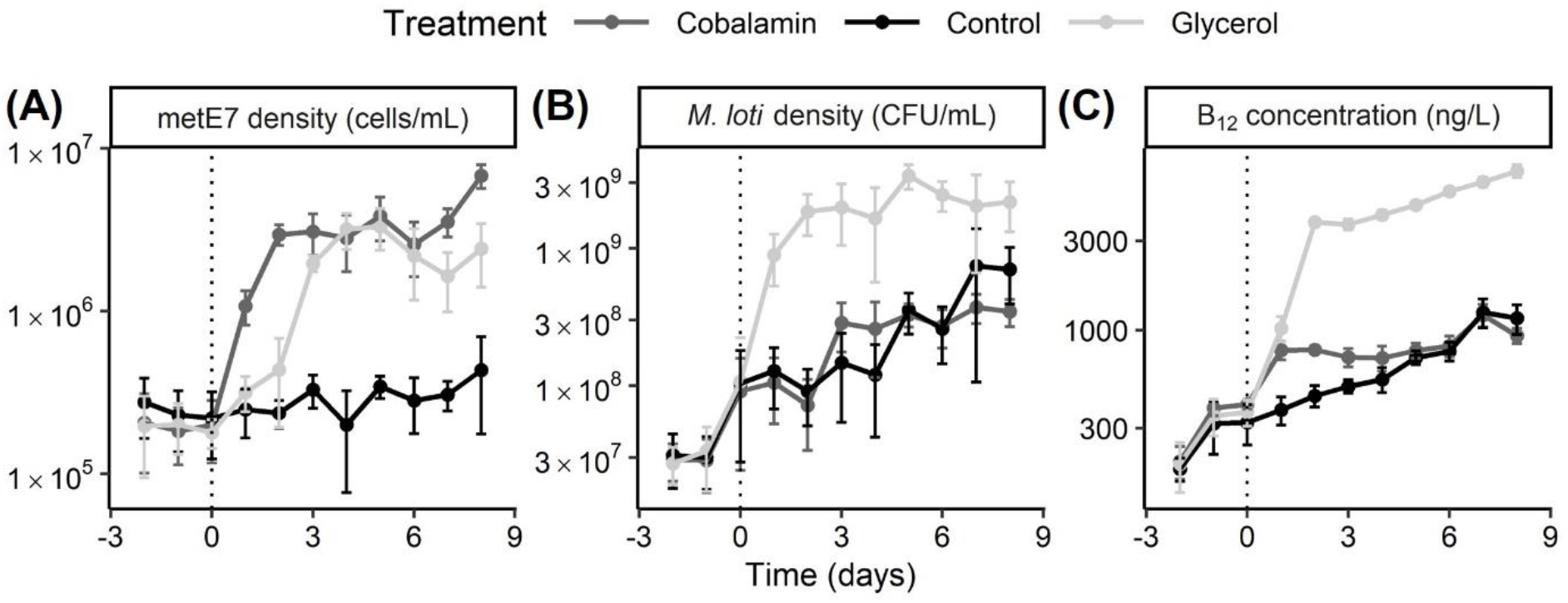
metE7 and *M. loti* only partially support each other’s nutrient requirements. *M. loti* and metE7 were cultured semi-continuously (10% volume replacement per day) under the same conditions as before, but with the addition of 200 ng·L^-1^ of B_12_, 0.02% glycerol or nothing from day 0. Periodic measurements of algal and bacterial density as well as B_12_ concentration in the cells and media were made **(A)** metE7 density increases in the B_12_ supplemented cultures, but also in the glycerol-supplemented cultures after a delay **(B)** *M. loti* density increases in response to glycerol but not B_12_ supplementation **(C)** Total B_12_ concentration increases more substantially in response to glycerol addition than B_12_ addition itself. Dark grey = 200 ng·L^-1^ of B_12_ addition, light grey = 0.02% glycerol addition, black = control. Error bars = standard deviation, n=4.

A similar study of the symbiosis between the B_12_-dependent alga *L. rostrata* and *M. loti* found that B_12_ production increased in the presence of *L. rostrata* (Kazamia *et al*., 2012; Grant *et al*., 2014). To determine whether the same was true with metE7 we collected further data from axenic cultures of *M. loti* grown in TP medium with glycerol or co-cultures of metE7 and *M. loti* in TP medium alone. Figure S6 reveals a clear positive correlation between B_12_ concentration and *M. loti* density, but that B_12_ produced per *M. loti* cell decreases at higher densities. Rather than produce more B_12_ when grown in co-culture with metE7, *M. loti* produced 45% less B_12_ in co-culture than axenic cultures of a similar density (p<0.0001). B_12_ in the media fraction increased less with increasing *M. loti* density in co-cultures than axenic cultures (p<0.0001), presumably due to metE7 B_12_ uptake. B_12_ levels in the cell fraction, conversely, were 50% higher in co-cultures (metE7 and *M. loti*) than axenic (*M. loti* alone) cultures (p<0.0001). Although we could not separate algal and bacterial fractions to measure cellular B_12_ in each, we hypothesised that the reduced B_12_ level in the media of co-cultures might lead to lower bacterial intracellular levels and hence could relieve suppression on B_12_ riboswitch-controlled B_12_ biosynthesis operons (Nahvi *et al*., 2004). We therefore tested the effect of removing B_12_ from the media of *M. loti* cultures either by replacing the media entirely or using metE7 to take up dissolved B_12_. B_12_-deprived metE7 was capable of absorbing, and so removing, most of the B_12_ released by *M. loti* in a manner that B_12_-saturated metE7 could not (Figure S7), and yet there was no subsequent increase in B_12_ production by the bacterial cells under that condition. Similarly, entirely refreshing the media, and so removing all B_12_ from the *M. loti* culture, did not increase B_12_ production. Therefore, it seems unlikely that *M. loti* responds to metE7 B_12_ uptake by increasing B_12_ synthesis.

### Greater bacterial B_12_ release increases both algal and bacterial growth in co-culture

It was not particularly surprising that metE7 did not increase *M. loti* B_12_ production, since these species may not encounter one another in the natural environment. Moreover, there would be no advantage to *M. loti* to provide more B_12_ to the naturally B_12_-independent *C. reinhardtii*. However, an evolved B_12_ dependent alga like metE7 might have a more passive and indirect way of increasing B_12_ in its environment: B_12_-dependent algae that happen to co-occur with higher B_12_ providers would grow faster, produce more photosynthate and so improve the growth of the B_12_ producers. To study the effect of B_12_ provision, we first compared the wild type strain of *M. loti* with a B_12_ uptake transporter (BtuF) mutant but found no significant difference in B_12_ production or release (Figure S8). The B_12_-producing rhizobial strains we had initially tested also did not have substantially different rates of B_12_ release, and due to their different growth rates and different physiologies, determining whether any effect on algal growth was due to B_12_ release would have been challenging. To address this we took advantage of two strains of *E. coli* engineered to produce B_12_. One of these strains lacked BtuF and our preliminary observations indicated that more B_12_ was found in the media of this strain. *E. coli*, a gut bacterium, does not naturally co-occur with *C. reinhardtii*, and most strains do not naturally produce B_12_, so we decided that these strains could be useful to test the importance of B_12_ release (not just B_12_ production) in supporting B_12_-dependent algae.

We cultured the two *E. coli* strains, ED656 and ΔbtuF, in TP media with 0.1% glycerol, under the same conditions as were used for *M. loti*, and then measured the cell density and B_12_ after four days. Figure 4A shows that ED656 and ΔbtuF both grew to a similar density of approximately 10^8^ CFU·mL^-1^, but ΔbtuF produced 50% more B_12_ (p<0.01) (Figure 4B).

**Figure 4.**
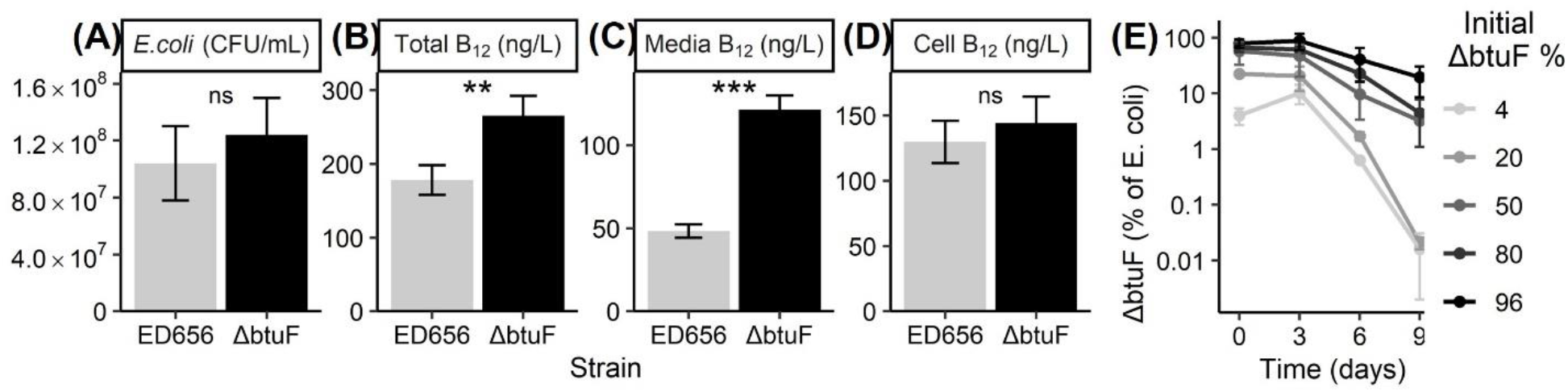
Growth and B_12_ production of two *E. coli* strains engineered to synthesise vitamin B_12_. ED656 expresses BtuF, a protein involved in B_12_ uptake, while ΔbtuF is a kanamycin resistant *btuF* knockout. *E. coli* cells were inoculated at 10^7^ cells·mL^-1^ and grown for 4 days in TP media with 0.1% glycerol (v/v), at 25 °C, and with illumination for a 16:8 hour light:dark period at 100 μE·m^2^·s^-1^. **(A)** *E. coli* density in colony forming units per mL. B_12_ measured by bioassay in the **(B)** whole culture, **(C)** supernatant after centrifugation at 10,000 g for 2 minutes, or **(D)** pellet after centrifugation. **(E)** *E. coli* strains were grown as above but were initially inoculated at different starting percentages: 4, 20, 50, 80, or 96% ΔbtuF with the remainder made up with ED656. Cultures were maintained by diluting 10,000-fold on day 3 and 6 after CFU density measurements. CFU density of ΔbtuF and both strains combined were measured over 9 days by plating a dilution series onto LB agar plates with or without kanamycin (50 μg·mL^-1^), respectively, and counting the colonies after an overnight incubation at 37°C. For panels A-D n=5, for panel E n=2 at each starting concentration of ΔbtuF.

Importantly, almost all this additional B_12_ was in the media fraction (Figure 4C), such that levels were 1.5-fold higher (p<0.001) for ΔbtuF than ED656, whereas the cellular B_12_ was not significantly different between *E. coli* strains (Figure 4D). Because the growth rates of ED656 and ΔbtuF were similar we designed a more sensitive experiment to distinguish them: after mixing both strains in different proportions the cultures were maintained and measured over nine days under the same conditions as described above with a 10,000-fold dilution on day 3 and 6. The proportion of ΔbtuF cells was substantially lower after 9 days than on day 0 irrespective of the starting concentration, indicating that ED656 had a faster growth rate (Figure 4E). Since the strains are otherwise isogenic this difference can be attributed to the lack of BtuF, the presence of the kanamycin resistance cassette, or both.

Due to the higher B_12_ production and release by ΔbtuF we predicted that it would support metE7 to a greater extent than ED656. Coculturing metE7 and either of the *E. coli* strains in TP media as before revealed that after 3 days metE7 cell density was over 100-fold greater (p=0.00014) in co-culture with ΔbtuF (Figure 5A). This was unlikely to be purely due to increased growth of ΔbtuF, however, because we found that although ΔbtuF grew to a greater extent than ED656, it was only by 1.6-fold (p=0.046) (Figure 5B). This resulted in the ratio of bacterial to algal cells for ED656:metE7 being much higher than the ΔbtuF:metE7, at 10,000:1 and 170:1 respectively (p=0.0023). When the *E. coli* strains were spotted out at equal densities on TP agar 10 mm away from metE7 and allowed to grow for 30 days, it was visibly clear that ΔbtuF supported metE7 to a greater extent than ED656 (Figure 5D).

**Figure 5.**
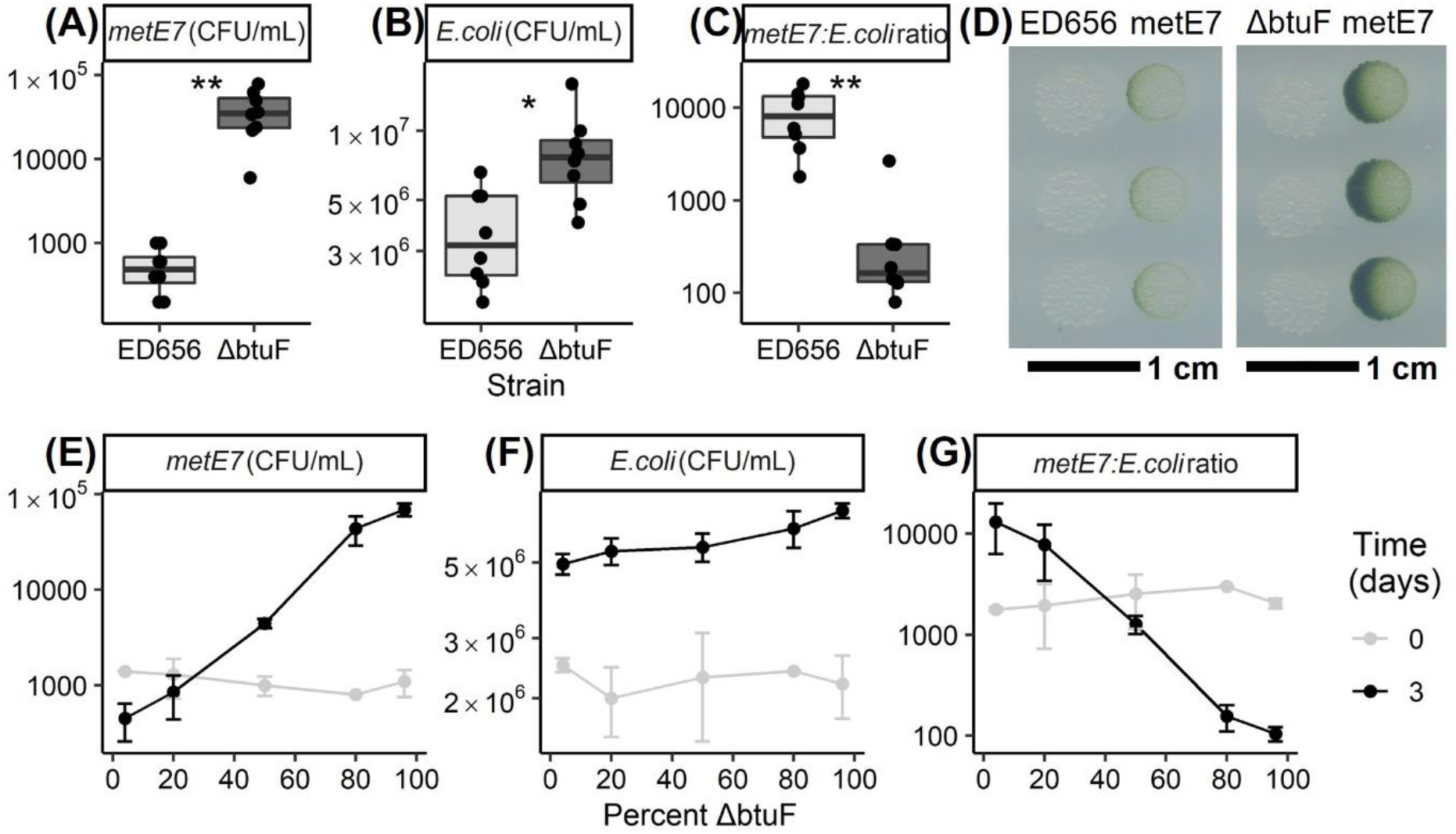
The ΔbtuF mutant of *E. coli* releases more B_12_ and is better than the isogenic parent ED656 at supporting metE7. For panels A, B, and C metE7 was co-cultured with either *E. coli* ED656 or ΔbtuF for three days at 25 °C with shaking at 120 rpm and constant illumination at 100 μE·m^2^·s^-1^. **(A)** metE7 colony forming unit (CFU) density on day three of co-culture with either *E. coli* strain. **(B)** ED656 or ΔbtuF CFU density on day three of co-culture with metE7. **(C)** Ratio of CFUs of *E. coli*:metE7 on day three of co-culture. **(D)** Photograph of 5 μL droplets of ED656 or ΔbtuF cultures at 10^5^ cells·mL^-1^ spotted 10 mm from metE7 cultures at 10^5^ cells·mL^-1^ on TP agar and incubated for 30 days at 25 °C with constant illumination at 100 μE·m^2^·s^-1^. For panels E, F, and G metE7 was co-cultured for three days under the same conditions as above with a mix of ΔbtuF and ED656 strains of *E. coli*, where the percentage of ΔbtuF in this mix was 4, 20, 50, 80 or 96%. **(E)** metE7 colony forming unit (CFU) density on day three of co-culture with different mixtures of *E. coli* strains. **(F)** Total *E. coli* (ED656 + ΔbtuF) CFU density on day three of co-culture with metE7. **(G)** Ratio of CFUs of *E. coli*:metE7 on day three of co-culture. For panels A-C n=7-8, and for E-F n=2 at each mixture.

To further investigate the difference in how well ED656 and ΔbtuF could support metE7, we initiated tricultures, all of which had the same density of metE7, but which had different ratios of ED656 to ΔbtuF, namely 1:24, 1:4, 1:1, 4:1, and 24:1. After three days in these tricultures it was clear that the higher the percent of ΔbtuF the higher the density that metE7 achieved (Figure 5E, p<0.0001). This was despite there being a slight negative correlation on day 0 in metE7 density vs percent of ΔbtuF (p=0.034). Furthermore, there was a smaller but still significant (p<0.0001) positive correlation between percentage ΔbtuF and total bacterial (ΔbtuF + ED656) growth (Figure 5F). These results suggest that not only do higher B_12_ producers and releasers better support a B_12_ dependent alga, but that they might also benefit themselves as a result of doing so.

## Discussion

In this study, we used an experimentally-evolved B_12_ dependent alga to ask how one that arose in the natural environment might survive when supported by B_12_-producing bacteria. We showed that the B_12_ dependent mutant of *C. reinhardtii*, metE7 (Helliwell et al. 2015), and the rhizobium *M. loti* could support one another’s growth, although metE7 grew more slowly than the B_12_-independent ancestral strain in coculture with *M. loti*, and B_12_ addition to the coculture increased metE7 growth. Although there are several reports of regulated interactions between algae and bacteria (Seyedsayamdost *et al*., 2011; Amin *et al*., 2015; Dao *et al*., 2020), we found no evidence that *M. loti* produced more B_12_ in response to encountering metE7. Nonetheless, we subsequently found that higher B_12_ providers supported more productive co-cultures, increasing both algal and bacterial growth. We therefore hypothesise that in heterogeneous environments B_12_ auxotrophs might colocalise with higher B_12_ providers due simply to positive feedback loops ensuring their relatively higher group fitness.

metE7 was previously shown to be B_12_-dependent (Helliwell *et al*., 2015, 2016), and *M. loti* known to produce enough B_12_ to support algal B_12_ auxotrophs (Kazamia *et al*., 2012; Helliwell *et al*., 2018; Peaudecerf *et al*., 2018), but here we more accurately determined the B_12_ requirements of metE7 grown under different trophic conditions, and revealed the dynamics of *M. loti* B_12_ production and release. metE7 had a significantly lower requirement for B_12_ (EC50∼10 ng·L^-1^) under phototrophic conditions than optimal mixotrophic conditions, presumably due to a slower growth rate and a lower carrying capacity (Fig S2). This requirement is similar to laboratory cultures and environmental samples of many fresh and saltwater B_12_-dependent algae, which mostly show half-saturation constants below 100 ng·L^-1^ (Sañudo-Wilhelmy *et al*., 2006; Tang *et al*., 2010; Helliwell, 2017). *M. loti* released sufficient B_12_ to raise the concentration in the media to almost 1000 ng·L^-1^ of B_12_ over six days of culture in media optimised for *C. reinhardtii*, which is considerably higher than is found in most aquatic or soil environments (Daisley, 1969; Sañudo-Wilhelmy *et al*., 2012; Barber-Lluch *et al*., 2021), although over 100,000 fold lower than is produced industrially by fermentation using *Propionibacterium shermanii* or *Pseudomonas denitrificans* (Acevedo-Rocha *et al*., 2019).

There is no known bacterial system for exporting corrinoids, but exogenous B_12_ addition has been shown to reduce bacterial B_12_ release, as has nutrient deprivation, suggesting some degree of regulation (Bonnet *et al*., 2010; Piwowarek *et al*., 2018). Therefore, B_12_ production per cell may have declined as *M. loti* entered nutrient-limited stationary phase (Figure S6) or because B_12_ accumulation caused negative feedback on B_12_ synthesis, although Figure S3 suggests this is unlikely as B_12_ addition did not affect B_12_ production. metE7 absorbed a large quantity of the B_12_ produced by *M. loti* from the medium over a short time period (absorption half-time of 4 minutes - see Figure 1C). Previous work has not quantified the dynamics of B_12_ uptake in *C. reinhardtii* at this resolution, but one study found that within one day *C. reinhardtii* cells absorbed 12,000 molecules per cell (Fumio Watanabe *et al*., 1991), considerably less than the roughly 300,000 molecules per cell we found were taken up within one hour. Studies in *Euglena gracilis* discovered uptake of 400,000 molecules of B_12_ per cell within the first minute, which might be explained by the fact that *E. gracilis* has a cell volume that is approximately 20 times greater than *C. reinhardtii (Shehata and Kempner, 1977; Sarhan et al., 1980; Craigie and Cavalier-Smith, 1982)*.

In co-culture with *M. loti*, metE7 grew less well than its ancestral line suggesting that its growth was B_12_-limited, and yet the growth of *M. loti* was not significantly lower with metE7 than the B_12_-independent lines for almost the entire growth period (Figure 2). One potential explanation may be that the B_12_-limited metE7 cells released (including through cell death) a greater amount of organic carbon or a different spectrum of compounds leading to a greater bacteria:algal ratio. In fact, increased cell size, cell death and an increase in starch and triacylglycerides were all found to occur in metE7 on B_12_ deprivation (Bunbury *et al*., 2020). Figure 3A clearly indicates that the addition of glycerol allows a large increase in *M. loti* growth and subsequently B_12_ production (Figure 3C), which in turn is the likely cause of the increase in metE7 cell density. Previous studies of algal-bacterial mutualisms have investigated the effect of nutrient addition to co-cultures and have produced a variety of results. Two studies of the bacterium *Dinoroseobacter shibae* co-cultured with different algae found that adding vitamins B_1_ and B_12_, which were required by the algae, actually improved bacterial growth by a greater amount (Cruz-López and Maske, 2016; Cooper *et al*., 2019). Our results were more similar to a study of *L. rostrata* and *M. loti*, which found that B_12_ addition improved algal growth with little to no effect on *M. loti* (Kazamia *et al*., 2012; Grant *et al*., 2014). Unlike the *L. rostrata* study, however, we found that after induction of *M. loti* growth by addition of glycerol, algal density subsequently increased, causing the bacterial:algal ratio to return towards pre-addition levels (Figure S5).

Across much of the eubacteria, there is a highly conserved regulatory riboswitch element that is found upstream of B_12_ biosynthesis and transport genes (Rodionov *et al*., 2003). In *M. loti* these elements are found upstream of B_12_ biosynthesis operons, and B_12_ suppresses the expression of some of these genes, likely resulting in reduced B_12_ production. We hypothesised that metE7 could compete for extracellular B_12_ and so indirectly decrease *M. loti* intracellular levels and potentially, therefore, increase B_12_ production. Although the media of dense co-cultures contained significantly less B_12_ than axenic cultures of *M. loti* with similar densities, this did not result in increased total B_12_ levels (Figure S6). The only fraction in which B_12_ was higher in co-culture was the cellular fraction, presumably because in the co-culture this includes both algal and bacterial cells. The short-term effects of perturbing *M. loti* cultures similarly did not indicate that either metE7 addition or B_12_ removal increased B_12_ production per *M. loti* cell (Figure S6). The only way that the addition of metE7 appeared to affect B_12_ production was through increased growth of *M. loti*, presumably through providing a small amount of photosynthate, and this occurred irrespective of whether metE7 absorbed any B_12_ from the media.

After finding that metE7 did not induce *M. loti* to synthesise more B_12_, in contrast to *L. rostrata* (Kazamia *et al*., 2012), it seemed unlikely that *C. reinhardtii* would have evolved any more complex measures to increase B_12_ supply such as partner selection or sanctioning (Leigh, 2010; Chomicki *et al*., 2020). However, we hypothesised that group selection might superficially resemble partner choice, because those bacteria that produced more B_12_ would increase algal growth and so benefit from increased photosynthate. A ΔbtuF mutant of *M. loti* did not release more B_12_ than the wild type (Figure S8) nor did *M. loti* appear to take up B_12_ (Figure S3). In another rhizobium, *S. meliloti*, it was found that the previously annotated *BtuCDF* genes were actually involved in cobalt uptake, not cobalamin uptake (Cheng *et al*., 2011), however cobalamin supplementation did appear to increase the growth of *S. meliloti (Campbell et al., 2006)*. If the homologs in *M. loti* also do not transport cobalamin, this would explain why ΔbtuF showed no difference in B_12_ release. More work will be necessary to test whether other rhizobia also do not take up B_12_, and whether the lack of B_12_ uptake predicts the ability to support B_12_ auxotrophs.

BtuF in *E. coli* does bind and transport B_12_ (Borths *et al*., 2002), and so we generated a B_12_-producing strain (ED656) of *E. coli*, and a *btuF* mutant (ΔbtuF) of this strain that did release more B_12_ (Figure 4). Furthermore, ΔbtuF was better able to support metE7 in co-culture, and subsequently grew better itself (Figure 5). Although ΔbtuF resulted in greater productivity in co-culture than the parental B_12_-producing *E. coli* strain (ED656), it also had a slower growth rate than the latter as indicated by its decreasing proportion in co-cultures with the parental strain (Figure 4E). This would suggest that in a homogeneous co-culture with a B_12_-dependent alga, ED656 would eventually dominate ΔbtuF and lead to the steady decline and possible collapse of the co-culture.

Generalising the example mentioned above to an ecological scenario where cooperative strains are at a disadvantage to ‘cheaters’ (those that contribute less or overexploit a public good) is frequently labelled as a ‘tragedy of the commons’ or a ‘prisoner’s dilemma’ (Hardin, 2009). These game theory concepts describe how it can be optimal at the group level for individuals to cooperate while simultaneously being optimal for each individual to cheat, such that the rational outcome (Nash equilibrium) is also the worst outcome overall (Nash, 1953). The fact that mutualistic interactions nevertheless abound in nature indicates that this dilemma is solvable, and spatial structure is often at the heart of these solutions (Stump *et al*., 2018). In multicellular hosts with microbial symbiotic partners, spatial structure is often manifested as compartmentalisation, allowing hosts to control, sanction, or reward their endosymbionts (Chomicki *et al*., 2020). In aquatic microbial symbioses, it is the phycosphere of separate algal cells that can provide spatial structure for the stable establishment of different bacterial communities (Kimbrel *et al*., 2019; Durán *et al*., 2021), and vertical transmission (which favours mutualism) (Crespi, 2001) of bacteria to algal daughter cells might even be more common than in plant-rhizobia symbioses.

In our example of an algal B_12_ auxotroph, whether the alga and its associated community figuratively sinks or swims is at least partially dependent upon whether those bacteria provide sufficient B_12_. It is therefore not unreasonable to hypothesise that bacteria that release more B_12_, such as the ΔbtuF mutant discussed here, could out-compete those that release less, if they co-occur with algal B_12_ auxotrophs such as metE7.

There may be similar advantages of enhanced B_12_-producers feeding B_12_-auxotrophs within further complex microbial communities, such as those found within the gastrointestinal microbiome, where B_12_ has been shown to be a key modulator of the ecosystem (Degnan, Taga, *et al*., 2014). Within the microbiome, better B_12_-producers may well be favoured partners of auxotrophs, but the situation is further complicated by the observation that bacteria produce a range of non-cobalamin corrinoids (Bryant *et al*., 2020), which may either help specific community formation or prevent predation. This complication is not found within bacterial-algal interactions since algae, like humans, appear to utilise just cobalamin (Helliwell *et al*., 2016).

In summary, our results suggest that B_12_-producing bacteria could support newly evolved algal B_12_ auxotrophs, but do not imply that they would drive the evolution of B_12_ auxotrophy. However, that is not to say that bacterial B_12_ production in the environment would be insufficient to do so. After all, in nature, purely bipartite, reciprocal relationships are probably rare, and heterotrophic bacteria and B_12_-dependent algae are likely to rely on multiple sources of organic carbon and B_12_, respectively. Furthermore, B_12_ auxotrophs may naturally have lower B_12_ requirements, or evolve lower requirements over time. Finally, natural selection tends to remove species that produce metabolites in excess of their own needs, but in a structured environment with photosynthetic B_12_ auxotrophs, we propose that it may be beneficial for a bacterium to produce and release more B_12_ rather than compete for its uptake.

## Experimental Procedures

### Strains

The wild-type *Chlamydomonas reinhardtii* strain used in this work was derived from strain 12 of wild type 137c. The two other mentioned strains, metE7 and revertant, were derived from this strain by experimental evolution: selection for rapid growth under 1000 ng·L^-1^ B_12_ produced a metE (B_12_-independent methionine synthase) mutant via insertion of a type II transposable element, which was called S-type (Helliwell *et al*., 2015). Subsequent excision of this transposon to repair the wild-type sequence of METE, produced the B_12_-independent revertant strain. In one instance, imprecise excision of the transposon left 9bp behind to produce a genetically stable B_12_-dependent strain, metE7 (Helliwell *et al*., 2015). *Mesorhizobium loti*, a soil-dwelling rhizobium that forms facultative symbiotic relationships with legumes, was used as a B_12_ producer in most co-culture experiments. Strains MAFF303099, the sequenced strain (Kaneko *et al*., 2000) and ΔbtuF and ΔbluB mutants from the STM library (Shimoda *et al*., 2008) were used. Two *Escherichia coli* strains, ED656 and ED662 (ΔbtuF), were also used as B_12_-producers. *E. coli* strain ED656 was constructed in a similar fashion to ED741 (Young *et al*., 2021). ED656 was generated from E. coli MG1655 by engineering it to contain a set of B_12_ biosynthesis genes. Strain ED656 is MG1655 with (Plac)-T7RNAP / (T7P)-cobA-I-G-J-F-M-K-L-H-B-W-N-S-T-Q-J-D-bluE-C-bluF-P-U-B-cbiW-V-E-btuR-R. All the genes were cloned individually in pET3a and then subcloned together using the “Link and Lock” method (Deery *et al*., 2012). ED662 (ΔbtuF) was derived from ED656 by insertion of a kanamycin resistance cassette in place of btuF. *E. coli* JW0154 ΔbtuF::KmR (Keio collection, *E. coli* Genetic Stock Center) was used for the construction of ED662 (= ED656 ΔbtuF::KmR) by P1-mediated transduction (Baba *et al*., 2006). *Salmonella typhimurium* AR3612, a metE and cysG double mutant, was used in bioassays to determine the concentration of B_12_ in samples (Raux et al. 1996). *Sinorhizobium meliloti* RM1021, *Rhizobium leguminosarum* bv. viciae 3841 and *Pseudomonas putida* were also initially assessed for B_12_ production and release.

### Algal and bacterial culture conditions

Algal colonies were maintained on Tris-acetate phosphate (TAP) (The Chlamydomonas Sourcebook) with Kropat’s trace elements except for selenium (Kropat et al. 2011) + 1000 ng·L^-1^ cyanocobalamin (CAS 68-19-9, Millipore-Sigma) agar (1.5%) in sealed transparent plastic tubes at room temperature and ambient light. Colony transfer by spreading on the agar surface using sterile loops was performed in a laminar flow hood. To initiate liquid cultures colonies were picked and inoculated into filter-capped cell culture flasks (NuncTM, ThermoFisher), or 24 well polystyrene plates (Corning®Costar® Merck) containing TAP or Tris minimal medium (The Chlamydomonas Sourcebook) at less than 60% volume capacity and supplemented with a range of B_12_ concentrations. Cultures were grown under continuous light or a light-dark period of 16hr-8hr, at 100 μE·m^-2^·s^-1^, at a temperature of 25 °C, with rotational shaking at 120 rpm in an incubator (InforsHTMultitron; Basel, Switzerland).

Bacteria were maintained in glycerol stocks stored at -80 °C. To initiate bacterial culture a small portion of the glycerol stock was removed by stabbing a pipette tip into the stock and spreading on LB and TY agar plates incubated at 37 °C overnight or 28 °C for 3 days for *E. coli* and *M. loti* respectively. After confirming the lack of contamination by colony morphology, single colonies were picked and inoculated into nunc flasks containing Tris minimal medium with 0.1% glycerol. Axenic bacterial cultures were incubated under the same conditions as algal cultures, as described above.

Co-cultures of algae and bacteria were initiated by inoculating them into Tris minimal medium from log-phase axenic cultures after dilution to ensure the optical density of algae and bacteria were roughly equal, except where specified. In some cases, co-cultures were acclimated for several days before making a dilution based on the algal cell density and starting measurements.

### Algal and bacterial growth measurements

Algal cell density and optical density at 730 nm were measured using a Z2 particle count analyser (Beckman Coulter Ltd.) with limits of 2.974 - 9.001 μm, and a FluoStar Optima (BMG labtech) or Thermo Spectronic UV1 spectrophotometer (ThermoFisher), respectively. Colony-forming units (CFU) of algal cells was determined by 10-fold serial dilution of a culture aliquot in growth media and spotting 10 μl volumes on TAP + 1000 ng·L^-1^ B_12_ 1.5% agar plates, followed by incubation at 25°C, 20 μE·m^-2^·s^-1^ of continuous light for 4 days and then counting the number of colonies at 100X magnification. *M. loti* and *E. coli* CFU densities were measured similarly but on TY or LB plates with incubation at 28 °C in the dark for 3 days or 37 °C overnight, respectively. *E. coli* ED662 (ΔbtuF) was distinguished from ED656 by its kanamycin resistance and hence its ability to grow on LB + 50 μg·mL^-1^ kanamycin plates. Single cells for all bacteria were too small to count with a light microscope. When bacteria were grown axenically, optical density was also used as a proxy for cell density with measurements at 600 nm on a FluoStar Optima (BMG labtech) spectrophotometer.

### Vitamin B_12_ quantification

A 1 mL aliquot of the culture to be tested was boiled for 5 minutes to release B_12_ into solution and then the growth response of a B_12_-dependent strain of *Salmonella typhimurium* (AR3612) incubated for 16 hours at 37°C in 50% 2*M9 media + 50% (diluted) boiled extract was quantified by measuring optical density at 600 nm (Raux et al. 1996). B_12_ concentration was calculated by comparing OD 600 nm of the cultures to a standard curve of known B_12_ concentrations using a fitted logistic model.

## Acknowledgements

We are grateful for helpful discussions with Dr Payam Mehrshahi and Dr Katrin Geisler (University of Cambridge) and Dr Katherine Helliwell (Marine Biological Association of the UK) and technical support from Geraldine Heath and Lorraine Archer. This work was supported by: the UK’s Biotechnology and Biological Sciences Research Council (BBSRC) Doctoral Training Partnership (grant no. BB/M011194/1) to F.B., A.S. and A.G.S.; BBSRC grant (BB/S002197/1) to M.J.W. and E.D.; the Cambridge Trusts (PhD scholarship to V.B.); the MELiSSA Foundation (FB European Space Agency) (grant no. CO-90-16-4078-02) to A.G.S. and E.H.; and Bill & Melinda Gates Foundation grant OPP1144 (PhD scholarship to A.H.).

## Author contributions

A.G.S and F.B. conceived and designed the research and drafted the manuscript. F.B., E.D. A.S, V.B. and E.L.H. participated in data acquisition and analysis. All authors helped draft and critically revised the manuscript and gave final approval for publication and agree to be held accountable for the work performed therein.

