## Supplemental Figures 1-8 for "Exploring the onset of B_12_-based mutualisms using a recently evolved Chlamydomonas auxotroph and B_12_-producing bacteria"

### Supplementary Figure 1

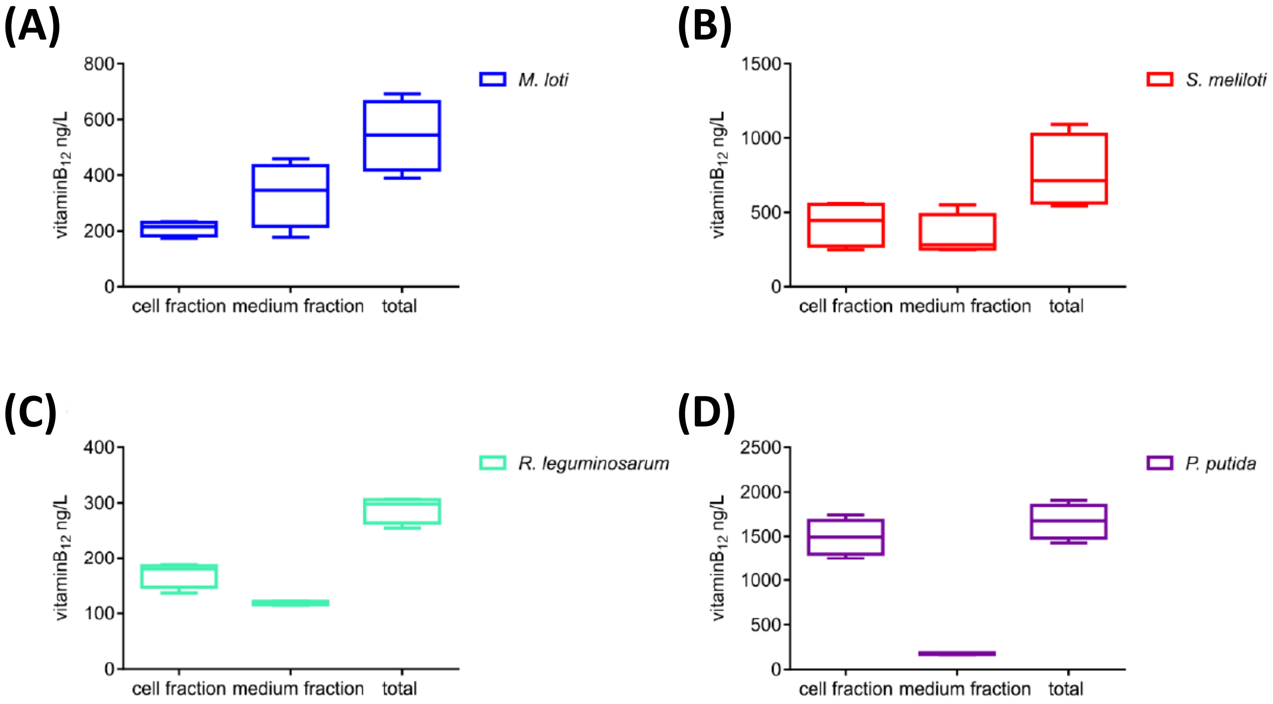


**Supplementary Figure 1.** Vitamin $B_{12}$ levels in the cell and media fraction of axenic cultures of four $B_{12}$-producing bacterial strains. Bacteria were grown in TP medium + 0.1% glycerol with illumination in a 12:12 hour light:dark period at 100 $\mu E \cdot m^{-2} \cdot s^{-1}$ and 25°C with rotational shaking at 120 rpm. After 6 days of growth, cultures were collected, centrifuged, and the pellet (cell fraction) and supernatant (medium fraction) separated and their $B_{12}$ content measured. The $B_{12}$ concentrations, in ng/L, are displayed as boxplots for **(A)** *Mesorhizobium. loti*, **(B)** *Sinorhizobium meliloti*, **(C)** *Rhizobium leguminosarum*, and **(D)** *Pseudomonas putida.* n=4 biological replicates.

### Supplementary Figure 2

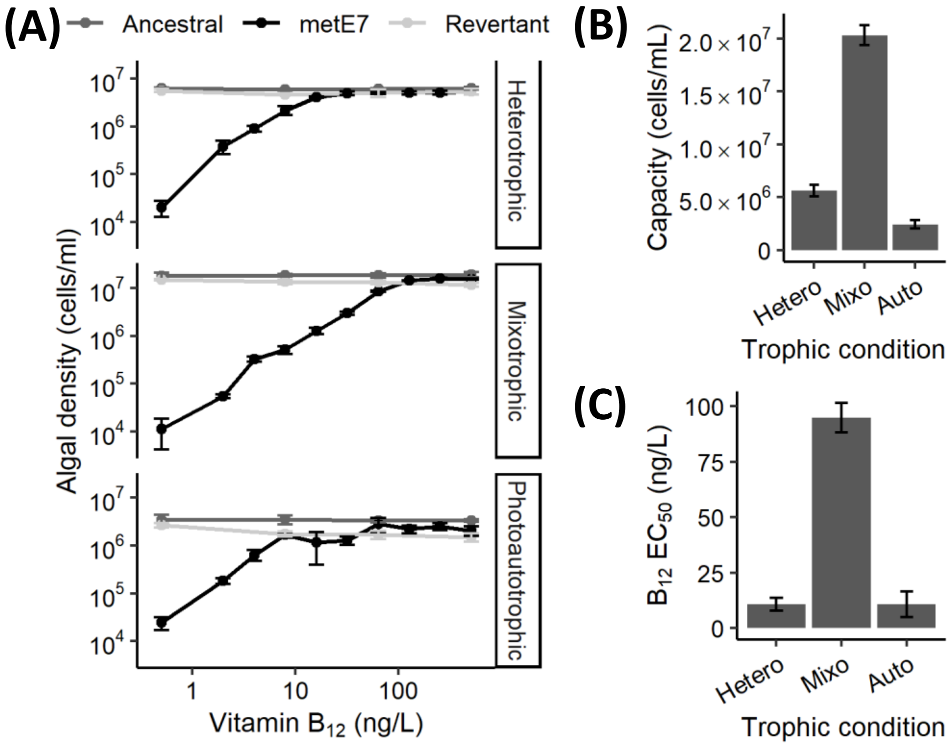


**Supplementary Figure 2.** Assessing the $B_{12}$ dependence of three lines of *C. reinhardtii* under different trophic conditions. The three lines include the 'ancestral' line prior to experimental evolution, 'metE7', a stable $B_{12}$-dependent line, and 'revertant', a $B_{12}$ independent line that had reverted from a $B_{12}$-dependent line. Cultures were grown heterotrophically (TAP medium in the dark), mixotrophically (TAP medium in continuous light), and photoautotrophically (Tris minimal medium in continuous light). $B_{12}$ concentrations ranged from 0.5 to 512 $ng{\cdot}L^{-1}$ and precultures of the algae, which were grown with 200 $ng{\cdot}L^{-1}$ $B_{12}$, were washed thrice and inoculated at a density of roughly 100 $cells{\cdot}mL^{-1}$. **(A)** Cell density was measured by particle counter after 6 days of growth for mixotrophic cultures or 8 days for heterotrophic and photoautotrophic conditions. **(B)** Estimated maximal density of metE7 at unlimiting $B_{12}$ concentrations calculated by fitting a Monod equation to data in panel A. **(C)** Estimated concentration of $B_{12}$ required to produce half the maximal density of metE7 cells under each trophic condition calculated by fitting a Monod equation to data in panel A. n=3-4, error bars = sd.

### Supplementary Figure 3

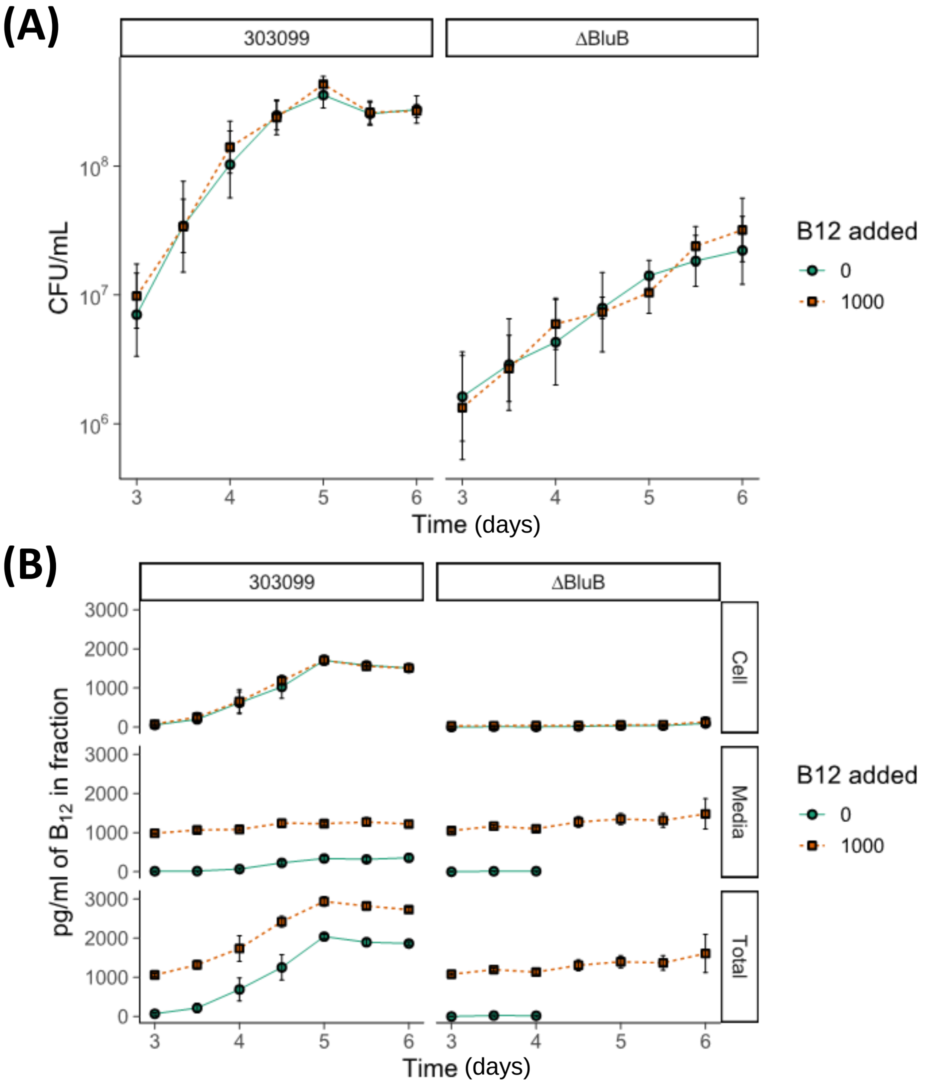


**Supplementary Figure 3.** Growth and $B_{12}$ uptake of *M. loti* strains. The wildtype (MAFF303099) and $B_{12}$ synthesis (ΔBluB) mutant were grown in Tris minimal medium supplemented with 0.1% glycerol at 100 $\mu E \cdot m^{-2} \cdot s^{-1}$, and at a temperature of 25$^{o}$C, with rotational shaking at 120 rpm over a period of 4 days (day 3-6) with (1000 $ng \cdot L^{-1}$) or without added $B_{12}$. **(A)** Viable cells (colony forming units) of *M. loti* MAFF303099 increased more quickly than the Δ BluB mutant, but there was no significant effect of $B_{12}$ on growth rate of either strain. **(B)** The addition of $B_{12}$ had no effect on the $B_{12}$ recovered in the cell fraction (top panel) indicating no $B_{12}$ uptake. Instead, all the added $B_{12}$ remained in the media (middle panel). Red lines = 1000 $ng \cdot L^{-1}$ of added $B_{12}$, Blue lines = no added $B_{12}$, Error bars = sd, n=4.

### Supplementary Figure 4

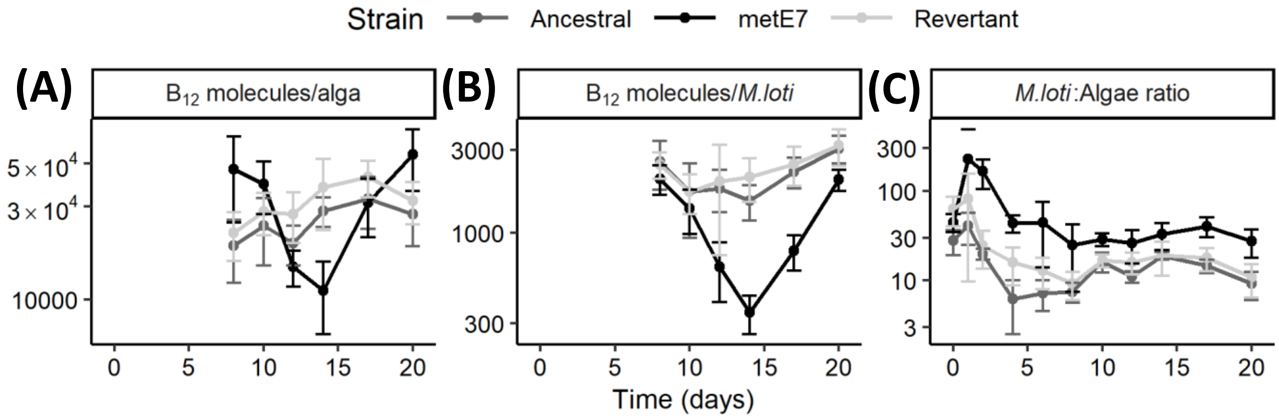


**Supplementary Figure 4.** Dynamics of the ratios of $B_{12}$, bacterial density and algal density during cocultures of *M. loti* and three strains of *C. reinhardtii* **(A)** $B_{12}$ concentration expressed as molecules of $B_{12}$ per algal cell reveal very similar levels although different dynamics for the three *C. reinhardtii* strains **(B)** $B_{12}$ concentration expressed as molecules of $B_{12}$ per *M. loti* cell reveal lower production in coculture with metE7 particularly around day 14 of coculture. **(C)** Bacteria:algae ratio was consistently higher in the metE7 coculture. Error bars = sd, n=5.

### Supplementary Figure 5

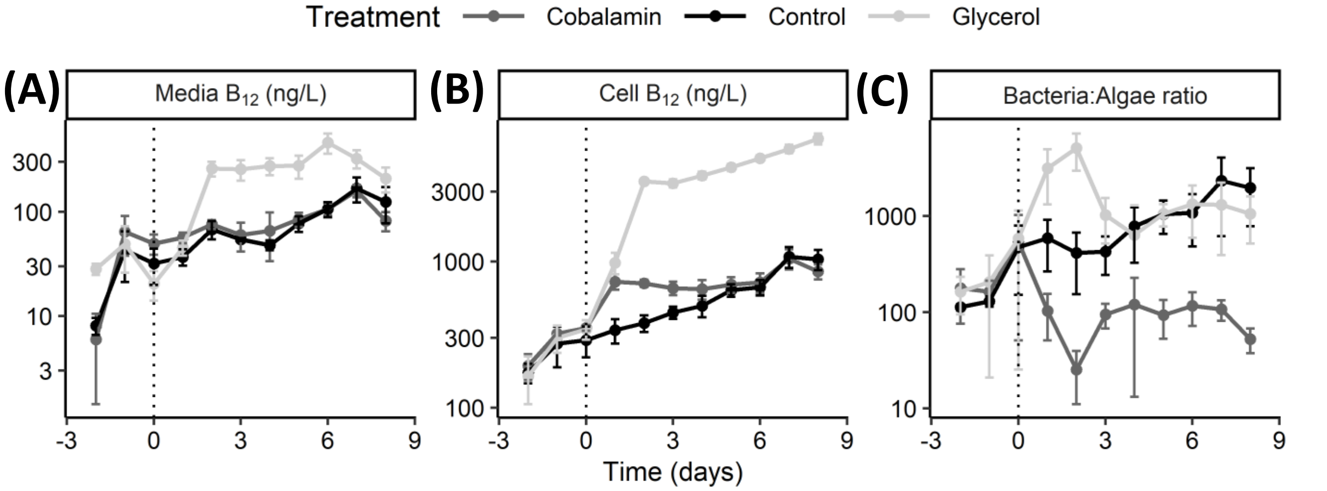


**Supplementary Figure 5.** Dynamics of $B_{12}$ concentrations in the cellular and media fractions and bacteria:algae ratio in metE7+*M. loti* cocultures perturbed by nutrient addition. **(A)** $B_{12}$ concentration in the media of cocultures reveals that the highest levels were found following addition of glycerol. **(B)** $B_{12}$ concentration in the cellular fraction reveals that glycerol addition caused significantly higher $B_{12}$ production. **(C)** Bacteria:algae ratio initially diverged after addition of glycerol or $B_{12}$ followed by a smaller convergence. Error bars = sd, n=4.

### Supplementary Figure 6

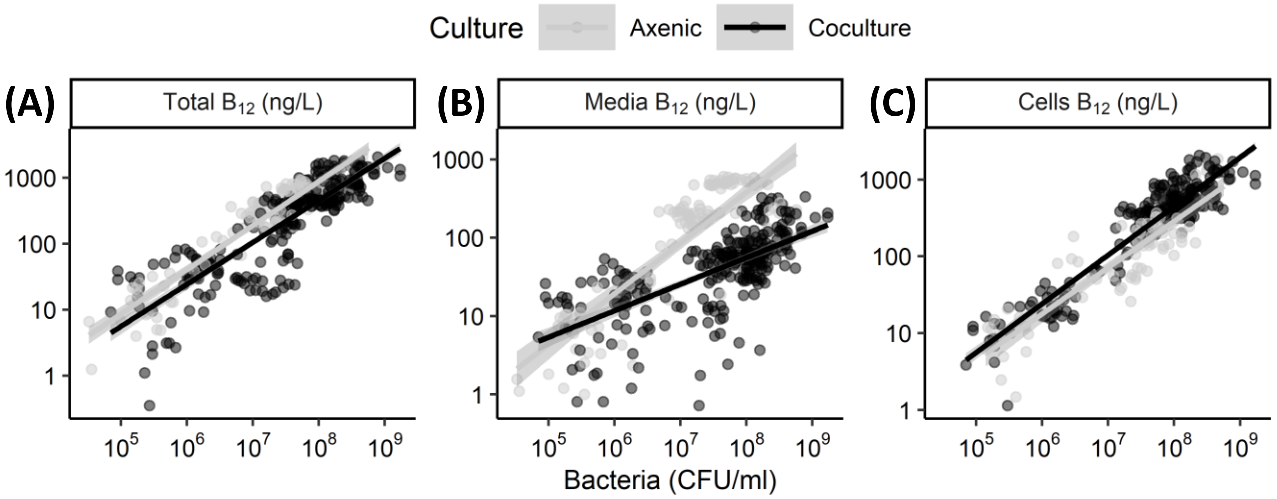


**Supplementary Figure 6.** *M. loti* does not increase $B_{12}$ production in the presence of metE7. Several axenic cultures of *M. loti* with supplemented glycerol and cocultures containing *M. loti* and metE7 (without glycerol) were grown in TP medium at 25°C with illumination at 100 $\mu E \cdot m^{-2} \cdot s^{-1}$ over a 16:8 hour light:dark cycle for up to 32 days or up until the cultures started to decline. $B_{12}$ measurements of the media and cell fraction were made periodically **(A)** Total $B_{12}$ is higher in axenic *M. loti* culture than coculture at the same *M. loti* density ($p<0.001$) **(B)** $B_{12}$ in the media is significantly lower in cocultures than axenic cultures at high *M. loti* densities ($p<0.001$). **(C)** Cellular $B_{12}$ is significantly higher in coculture than axenic culture at the same *M. loti* densities ($p<0.001$). Grey = *M. loti* axenic culture, black=metE7 + *M. loti* coculture. N(axenic)=106, N(coculture)=284, grey shaded region = 95% confidence interval.

### Supplementary Figure 7

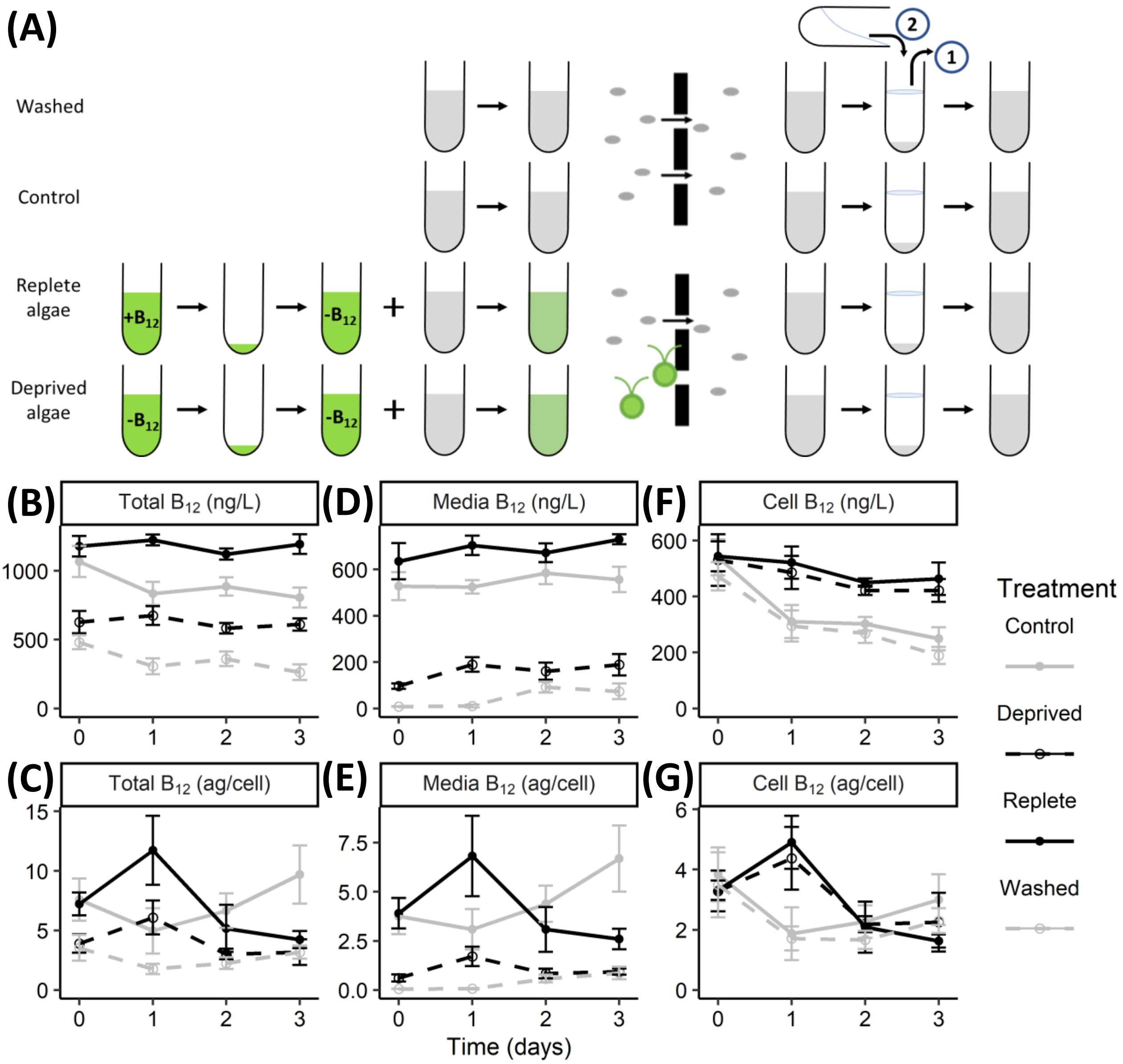


**Supplementary Figure 7.** $B_{12}$ production by *M. loti* following removal of $B_{12}$ from the culture media. **(A)** Experimental setup: Two sets of axenic *M. loti* cultures (grey) were inoculated with metE7 cells that were either saturated with (black solid) or starved of (black dashed) $B_{12}$ and incubated for 1 hour. All 4 cultures were then passed through a 5 µm filter, removing all metE7 cells but not *M. loti*. These *M. loti* cultures were centrifuged, and the supernatant replaced with fresh Tris-min media in treatment 'washed' (grey dashed), or otherwise resuspended without replacing the supernatant (grey solid). The resuspended, newly axenic *M. loti* cultures were grown for 3 days with illumination in a 16:8 hour period at 100 µE·m$^{-2}$·s$^{-1}$ and 25°C with rotational shaking at 120 rpm. **(B)** Total $B_{12}$ concentration in the culture, and **(C)** Total $B_{12}$ per *M. loti* cell. **(D)** $B_{12}$ concentration in the supernatant (media) after centrifuging an aliquot of the sample, and **(E)** media $B_{12}$ per *M. loti* cell. **(F)** $B_{12}$ concentration in the cell pellet after centrifuging an aliquot of the sample, and **(G)** cell $B_{12}$ per *M. loti* cell. Error bars = sd, n=4.

### Supplementary Figure 8

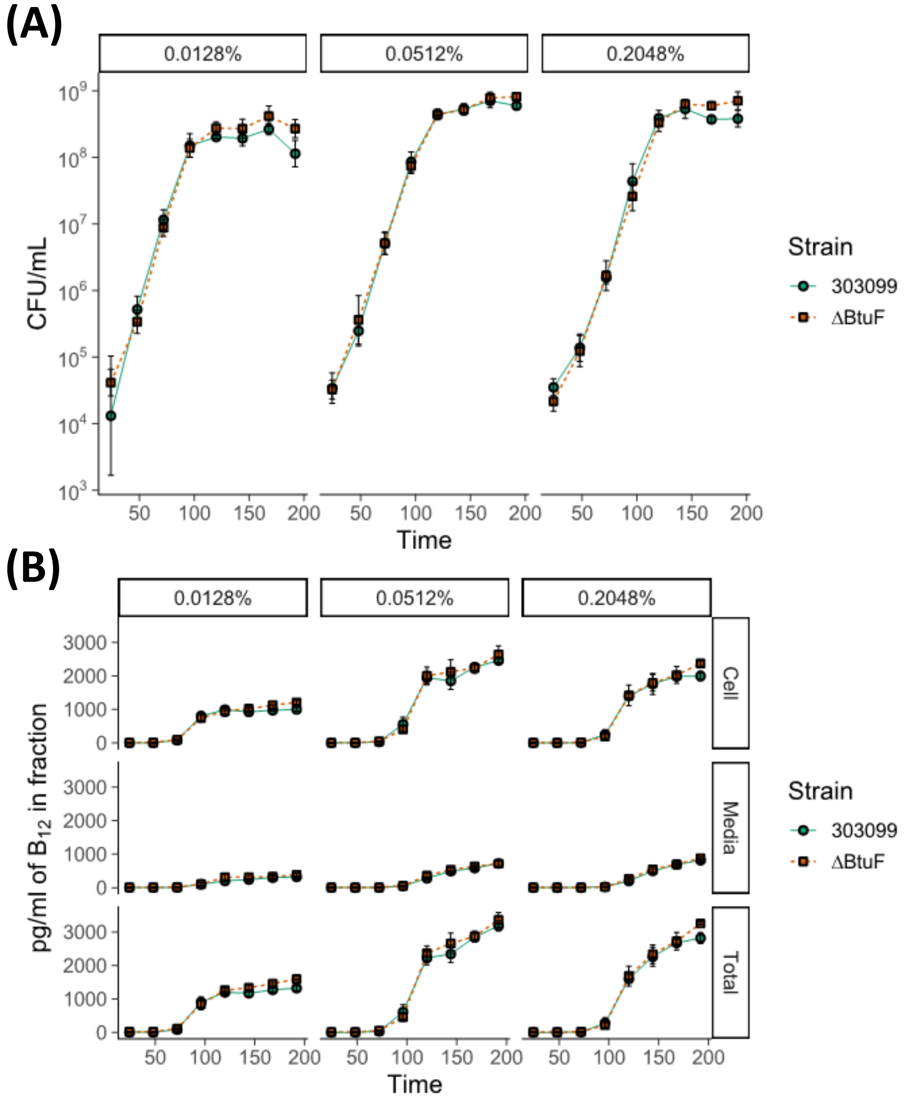


**Supplementary Figure 8.** Growth and $B_{12}$ release of *M. loti* strains. The wildtype (MAFF303099) and $B_{12}$ transporter (ΔBtuF) mutant were grown in Tris minimal medium supplemented with various concentrations of glycerol (indicated as a v/v percentage in the top panels) at 100 $\mu E \cdot m^{-2} \cdot s^{-1}$, and at a temperature of 25°C with rotational shaking at 120 rpm over a period of 8 days. **(A)** Viable cells (colony forming units) of *M. loti* 303099 increased over time at the same rate as the BtuF mutant and both strains showed improved growth on increasing the glycerol concentration from 0.0128% (v/v) to 0.0512%, but not with a higher concentration. **(B)** The amount of $B_{12}$ produced in the cells (top panel) and released into the media (middle panel) were not significantly different in the two strains, but as with the cell growth, did increase with the two higher glycerol concentrations. Blue lines = wildtype (MAFF303099), Red lines = ΔBtuF mutant, Error bars = sd, n=4.
